# Carbohydrate Hydrolytic Activity, Antibiotic Resistance and Stress Tolerance of *Lacticaseibacillus paracasei* BCRC-16100 and *Lacticaseibacillus paracasei* ZFM54 for Probiotics Using Genomic and Biochemical Approaches

**DOI:** 10.1101/2024.03.31.587454

**Authors:** Peter James Icalia Gann, Jimmbeth Zenila P. Fabia, Althea Gay B. Pagurayan, Ma. Joy Theresa Agcaoili, Ryan James J. Pascual, Suerte M. Baranda, Arc Josam J. Racho, Marvielyn P. Olivar, Jayson F. Cariaga, Alvin Domingo, Dionisio S. Bucao, Shirley C. Agrupis

## Abstract

Probiotics are microorganisms infused in products for health benefits including acceleration of nutrient digestion, however, it is also important to ensure the safety prior to incorporation. Here, we present evidence of the ability of two probiotic isolates, *Lacticaseibacillus paracasei* BCRC-16100 and *Lacticaseibacillus paracasei* ZFM54, in the (1) enhancement of carbohydrate digestion, (2) tolerance to stress, and (3) antibiotic resistance. Approaches include whole genome sequence (WGS) analysis and bioactivity assays. WGS revealed genes suggesting the ability of the two isolates to promote carbohydrate digestion, tolerance to stress and antibiotic resistance. Carbohydrate digestive ability was confirmed through a biochemical assay where the two isolates cause glucose release from starch. The two isolates also showed versatility in a range of temperature and alcohol concentration. For antibiotic resistance particularly on vancomycin, there are three mechanisms namely transporter control, transcriptional regulation, and efflux pump. Furthermore, promoter and transposable element analysis showed that some of the active antibiotic resistant (AbR) genes can be laterally transferred. Altogether, we show the potential of two probiotic isolates to be incorporated in products for improved carbohydrate digestion and the need to address the removal of active and mobile antibiotic resistance genes that may compromise safety.

## 1. INTRODUCTION

In the last ten years, there has been a significant progress in probiotic research with many studies suggesting the essential role of probiotics in maintaining human health. Probiotics is increasingly gaining a huge popularity and share in the market due to potential health benefits such as improved bioavailability of macronutrients, changing the diversity of the gut microbiota and treatment of several diseases. Microbes under the genera *Pediococcus, Lactococcus, Enterococcus, Streptococcus, Propionibacterium*, and *Bacillus* are generally considered as probiotics provided that they qualify screenings for function and safety [1-2].

Several studies have shown that probiotics can improve digestion of carbohydrates such as lactose [3] where lactose intolerant patients with supplemented with *lactobacilli* or *bifidobacteria* in their diet showed improved bioavailability and digestion of lactose [4-7]. Moreover, probiotics can also facilitate digestion of resistant starch such as fiber. When fiber passes through the stomach and small intestine undigested, the presence of *Bifidobacteria* in the large intestine can catalyze its hydrolysis [8-10]. There are also evidences on the enhanced protein digestion and absorption with probiotics which includes multiple mechanisms such as production of proteases, regulation of microflora to favor microbes with peptidase activities, and improved absorption of peptides and amino acids through accelerated transport in the intestine. The feasibility of lactobacilli derived proteases have been shown to catalyze hydrolysis of complex polypeptide chains [11]. Some of the most common protein hydrolytic enzymes in microbes are aspartic proteases, cysteine proteases, metalloproteases, and serine proteases [12]. These findings indicate the potential of probiotics to facilitate utilization of nutrients from food sources with low digestibility which consequently improves digestion and overall nutritional health.

In addition to the digestive benefits, probiotics can also alter the gut microbiome favoring groups of microbial flora that can improve health in three ways including (1) modifying the internal host environment, (2) change in the composition of the gut microbiome, and (3) inhibition of pathogen proliferation. First, a shift in the diversity of the gut microflora also changes the host gut internal environment through increase in the amount of specific metabolic compounds. Metabolites from probiotics including organic acids, short-chain fatty acids, teichoic acids, peptidoglycans, plasmalogens, neurotransmitters, biosurfactants, amino acids, and flavonoid-derived compounds such as desaminotyrosine, equol daidzein, noratirriol, terpenoids, and phenolic compounds has been found to cause positive effects on consumer health [13-15]. Second, observation animal models suggests that composition of the gut microbiome changes with probiotic supplementation [16]. A wide range of studies have shown that a perturbed microbial community in the gut can be restored using probiotics as it has been associated with competitive exclusion of pathogen binding [17-19]. This ability of lactic acid bacteria in modulating the gut microbiota is due to the production of antimicrobial peptides (AMPs) such as bacteriocins which function as a natural bacterial immune defense system that can be combined with the corresponding receptor of both narrow and broad spectrum microbiota, and other metabolites such as lactic acid, which are harmful to several pathogenic strains of microorganisms [20-23], thus, bacteria which produce lactic acid as end product of carbohydrate fermentation are generally recognized as safe (GRAS) by Food and Drug Administration (FDA). Scant nutrients in the environment trigger the production of bacteriocins for competition of space and resources, excluding potential pathogens in gut microbiome, thus altering the composition of intestinal microbiota. Lastly, alteration of the gut microbiome and modification of the host internal environment with the consumption of probiotics consequently treats gastrointestinal diseases in controlled clinical trials [24-25] and improves disease susceptibility [26].

Probiotics may provide multiple health benefits, but it is equally essential that microbes used for these purposes are non-pathogenic, non-toxic, and versatile to remain stable and viable for long periods of storage and harsh conditions for incorporation in both functional food and dietary supplements and drugs [20], and generally safe for the intended use and meets standards for purity, identity, and potency. Scrutiny of pathogenicity, physiological and metabolic activities, and intrinsic properties are required. Comprehensive assessment of probiotic strain intended to be utilized in foods and human supplements, despite the variation in regulatory requirements in each country must be highly considered [27] through ways including (1) assessment of physiological activities, (2) strain identity, and (3) antibiotic resistance. First, the industrial production of probiotic strains should be based on the properties of the strains involved and their ability to withstand stress during processing and storage [28]. Second, unambiguous identification of the probiotic strains of interest should be considered and whole genome sequence (WGS) is used for this purpose. Lastly, only probiotics strains which do not have antibiotic resistant (AbR) genes should be selected for use in food and supplements [29]. Phenotypic assessment of antibiotic resistant genes comprised of testing the probiotic strain to a set of clinically important antibiotics, and determination of colony forming units for each antibiotic exposure with respect to the original log of the strain culture. Annotated genome sequence of lactic acid bacteria is utilized to inspect genetic elements responsible for metabolic activities, physiological activities, hemolytic activity, toxins, and antibiotic resistance. The horizontal transfer capability of antibiotic resistance is further determined by gene identification either on a plasmid or in near proximity to mobile elements such as to transposases. The vancomycin-resistance phenotype present in several strains of *Lactobacillus* is one of the most used examples of intrinsic AbR which is attributed to properties of the cell wall preventing the binding of the antibiotic [30]. Based on EFSA in 2012, any functional antibiotic resistance genes found within the genome of probiotic strain should be characterized as intrinsic or transmissible, which will further determine the risk of potential spread, which is a major public-health concern nowadays [31]. Countries with modern regulatory structures utilize advanced technology in establishing probiotic safety profiles, with greater emphasis on whole genome sequencing, and relying less on *in-vivo* testing.

Candidate probiotic strains are ubiquitous in nature. They emanate from human and animal origins such as gastrointestinal tract and breast milk, various food biotopes like fermented food products and dairy products, as well as from different parts of plants [32]. Abundant sources of potential probiotic *Lactobacillus* strains were isolated from fermented food product of plant origin [33]. This broad range of inexhaustible sources provide a challenge for the lactic acid bacteria strains to adapt to various environments, and these abilities vary significantly among species and even at a strain level [34], hence a significant variation in their genomic and metabolomic profiles as well. This variation arises through processes such as genetic mutation, horizontal gene transfer, leading to variations in traits like phenotypic, metabolism, and antibiotic resistance. Additionally, environmental factors such as but not limited to nutrient availability, temperature, pH, and competition with other microorganisms exert selective pressures shaping bacterial populations, further contributing to the diversity in bacterial characteristics across various sources [35].

It is evident that probiotics is becoming increasingly popular to consumers as it offers multiple health benefits. This trend brings in the need to isolate probiotics from different sources, ensure safety and evaluate functional use when used for supplementation. Here, we elucidate the potential benefit of two probiotic isolates from nipa sap in improving digestion of carbohydrates and proteins. Moreover, we explain bottlenecks on safety with emphasis on antibiotic resistance and the underlying biomolecular mechanisms. Finally, we provide insights to work around the known limitations for safety of the two isolates.

## 2. MATERIALS AND METHODS

### 2.1. Isolation and Screening of Lactic Acid Bacteria (LAB)

The de Man Rogosa Sharpe agar (Sigma-Aldrich Canada Co.) supplemented with 1% CaCO3 was used as a selective medium for the cultivation and isolation of the two probiotic isolates from nipa sap. One hundred microliter of the serially diluted sample was aseptically inoculated in MRSA plate using spread plate method. The plates were incubated for 24h at 37°C. Individual colonies with colony morphology of a putative lactic acid bacterium grown overnight was further subjected to standard purification of bacterial cultures. Purified putative LAB undergone biochemical test consists of staining test using gram staining solutions (Sigma-Aldrich 111885) and catalase test using 3% H2O2. Moreover, hemolysin test using Sheep Blood Agar (Remel TSA w/ 5% Sheep Blood) was conducted with gram positive and catalase negative putative LAB. Isolates which exhibited γ-hemolysis were stored and sent for molecular identification.

### 2.2. Identification of Isolates by Capillary Sequencing

Genomic DNA (gDNA) of S1 and S2 was extracted using the Quick-DNA fungal/Bacterial Miniprep Kit (Zymo Research, USA) according to manufacturer’s protocol. PCR amplicons were subjected to purification using AMPure XP beads (Cat. No. 163881). One microliter of the purified PCR amplicons was loaded into 1% agarose gel run at 120 V for 45 min, with Invitrogen 1kb Plus DNA Ladder. Capillary sequencing involved the incorporation of fluorescently labeled chain terminator ddNTPs. The reaction components include amplicons, corresponding primers, and the ABI BigDye® Terminator v3.1 Cycle Sequencing Kit (Cat No. 4337455). The cycling parameters on the thermal cycler were as follows: pre-hold at 4°C; 96°C for 1min; 25 cycles at 96°C for 10s, 50°C for 5s, 62°C for 4min; and hold at 4°C. Ethanol precipitation was performed to remove unincorporated ddNTPs, excess primers, and primer dimers. Capillary electrophoresis was carried out on the ABI 3730xl DNA Analyzer using a 50cm 96-capillary array, POP7 Polymer (Cat No. 4393714), and 3730xl Data Collection Software v3.1. Base calling was performed using the Sequencing Analysis Software v5.4.

### 2.3. Whole Genome Sequencing

Library preparation was performed using the TruSeq DNA Nano Kit (Illumina, USA), and sequencing was conducted using an Illumina MiSeq instrument and a paired end read format of 2x150bp for 300 cycles at the Philippine Genome Center, Quezon City, Philippines.

### 2.4. Antibiotic Susceptibility Test

The probiotic isolates were subjected to antibiotic susceptibility using commonly used antibiotics (BD BBL Sensi-Disc) from various classes with various pharmacological actions, namely, natural penicillins, glycopeptides, aminoglycosides, macrolides, lincosamides, and fluorquinolones. Kirby-Bauer technique (disc-diffusion) was employed where one hundred microliters of bacterial culture was inoculated in the plate containing sterile Mueller Hinton Agar (TM Media, India) and discs of Vancomycin (30µg), Clindamycin (2µg), Gentamycin (10µg), Ofloxacin (5µg), Erythromycin (15µg), and Streptomycin (300µg) were used as antibiotic wafers. Zone of Inhibition (ZOI) was measured after 48h using digital caliper. The Results were interpreted as follows: resistant/R (<15mm), intermediate/I (16-20mm), and sensitive/S (>21mm). This test was done in three (3) replicates, and antibiotic-free disc plates were used as negative control. Moreover, viable cells of the probiotic isolates suspended in Mueller Hinton Broth (TM Media, India) containing the antibiotic with known concentration was quantified to correlate the semi-quantitative data obtained from disc-diffusion method using standard formula.

### 2.5. Carbohydrate Hydrolysis

Hydrolysis of *Lacticaseibacillus paracasei* BCRC-16100 and *Lacticaseibacillus paracasei* ZFM54 were qualitatively identified through agar well diffusion assay using a solution of 23g nutrient agar (TM Media, India) and 10g potato starch in 1L distilled H_2_O. 300μL of de Man Rogosa Sharpe broth (Sigma-Aldrich Canada Co.) were placed in 3 agar wells as control. 100μL aliquot of 48h bacterial cultures (1 – 2 × 10^7 cells approximately) were inoculated onto wells of the starch agar, with pH 6.5. The solidified agars were bored using sterile cork borer with a diameter of +10 mm. The plates were incubated separately for 24h and 48h at 35°C. The clearance zone was measured in mm for 24h and 48h using digital caliper (Linear*Tools*). The clearance indicates the carbohydrate hydrolysis activity of the two probiotic isolates.

### 2.6. Prediction of Promoter Elements

Bioinformatical tools such as BPROM and PRODORIC were used to predict the promoter elements involved in the expression of genes associated with antibiotic resistance, stress tolerance and hydrolytic activity of *Lacticaseibacillus paracasei* BCRC-16100 and *Lacticaseibacillus paracasei* ZFM54. The upstream regions of these genes were extracted from their whole genome sequences (WGS) and subjected to analysis in the BPROM website where results show possible –10 and –35 boxes of predicted promoters, their positions in the submitted sequence along with possible transcription factors. These transcription factors were further analyzed in the PRODORIC website’s virtual footprint in which their potential binding site with the highest relative score is shown in **Table S10**.

### 2.7. Gene Expression and Polymerase Chain Reaction

For gene expression, total RNA was isolated using Trizol (Invitrogen Inc.) and quantified using Nano-drop 2000 (Thermo-Fisher Inc). Two micrograms of total RNA were treated with RQ1-RNAse free DNase (Thermofisher Inc.), and one microgram of the DNase-treated RNA was used for cDNA synthesis using PrimeScript RT reagent kit (Takara Bio, CA, USA). The expression analysis was performed using TB green Premix Ex Taq II (Takara Bio, CA, USA) on Bio-Rad CFX 96 C1000 with following conditions: 95°C for 30s and 40 cycles of 95°C for 5s + 60°C for 30s. The product specificity was verified by the melt curve analysis. The Ct values of all the genes were normalized against *16s rRNA* fused protein as the reference gene. Primer sequences used for gene expression are in **Table S1**. The polymerase chain reaction (PCR) was performed on genomic DNA to amplify *16s rRNA* using Emerald Amp MAX PCR Master Mix (Takara Bio) using the primers presented in **Table S1**.

### 2.8. Data Analysis

Experiments subjected to statistical analysis were replicated three times. For bioactivity on macronutrient hydrolysis and other phenotypic characteristics between the two isolates, means were compared using studentized t-Test. Antibiotic resistance assays experiments were laid out in completely randomized design. Data were transformed with arcsine and was subjected to one-way ANOVA. Differences in the mean antibiotic resistance in different antibacterial drugs were determined using Tukey’s multiple comparison test. Statistical analyses were performed in SAS statistical software (version 9.4, SAS Institute Inc.) and results are presented in **Tables S2-9**.

## 3. RESULTS

### 3.1. Phenotypic and Genomic Characteristics and Identification of Isolates

There were two lactic acid bacteria isolated from nipa sap which were both subjected to phenotypic characterization and identification (**Figure 1a-c**). Both bacterial isolates were gram-positive with the shape of a bacilli. The two isolates showed negative reaction to 3% H2O2 and exhibited γ hemolysis as there is no formation of clear, greenish, or opaque zones around the colonies (**Figure 1a**). These phenotypes have been identified to be common in lactic acid bacterial isolates [36].

**Figure 1.**
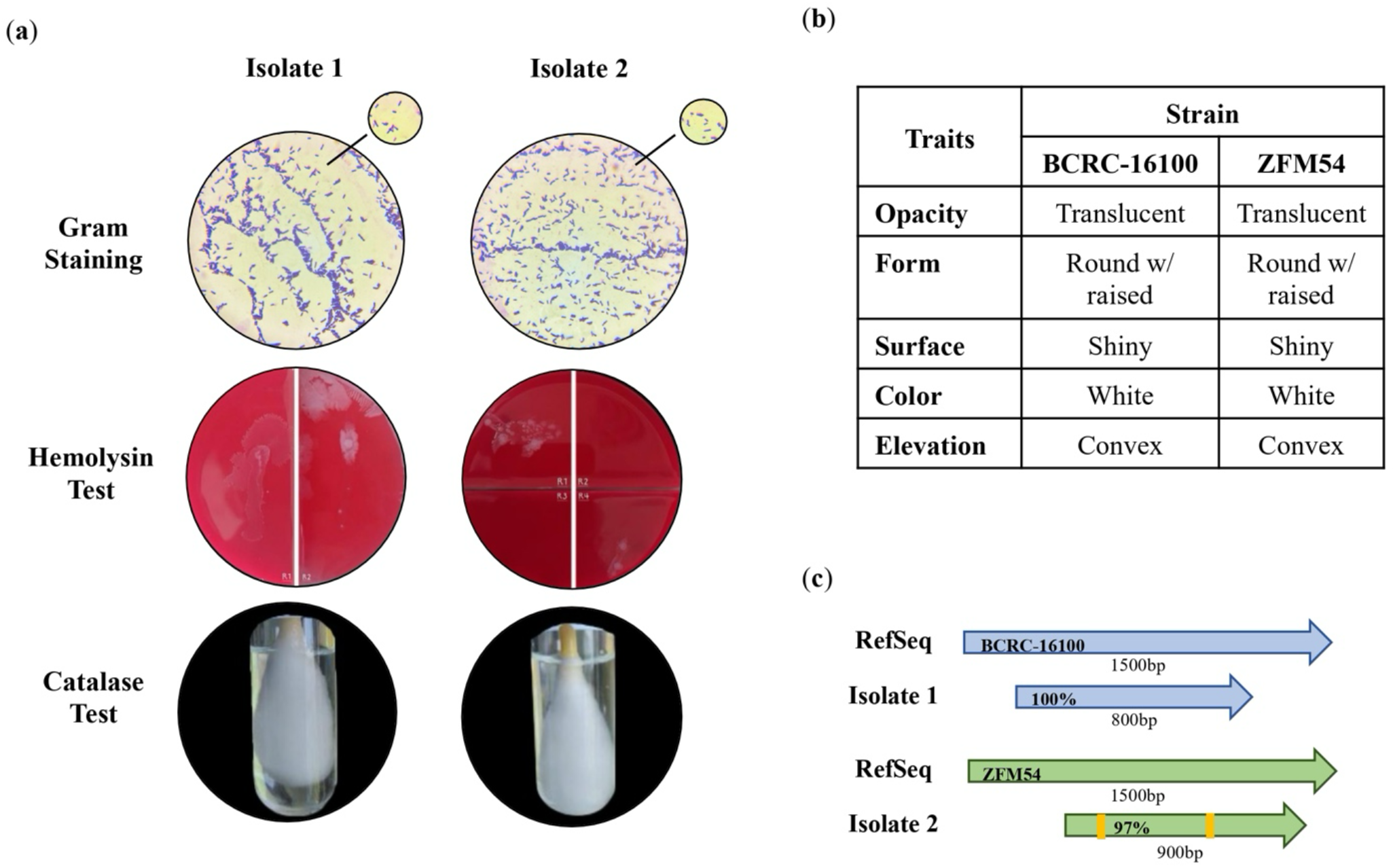
Phenotypic characteristics and identification of lactic acid bacterial (LAB) isolates from nipa sap, *Lacticaseibacillus paracasei* BCRC-16100 and *Lacticaseibacillus paracasei* ZFM54. (**a**) Tests for safety in the two isolates indicating gram positive, hemolysin negative, catalase negative for both, (**b**) observable physical appearance and (**c**) identity using 16s rRNA. Gram stain test (Purple cell morphology = gram positive, pink cell morphology = gram negative); hemolysin test (yellow to translucent inhibition = β-hemolysis, translucent with greenish inhibition = α-hemolysis, no clearing zones = γ-hemolysis; catalase test (presence of bubbles = positive, absence of bubbles – negative).

Through this, it is inferred that these putative lactic acid bacteria are safe for further utilization as probiotics that can be incorporated in drugs or functional food. The colonies of the two isolates were translucent, round, shiny and convex (**Figure 1b**) which are all consistent to the characteristics of a lactic acid bacteria described from other studies [37-42]. Using 16s rRNA, the isolates were identified as *Lacticaseibacillus paracasei* BCRC-16100 and *Lacticaseibacillus paracasei* ZFM54 with 100% homologies from the database sequences with accession numbers NZ_CP086132 and NZ_CP032637, respectively (**Figure 1c**). Despite genotypic similarities, alignment of genome sequences showed variability among the isolates and database hits (**Table S11**). Visualization of the whole genome of *Lacticaseibacillus paracasei* BCRC-16100 shows 2,885 contigs (3,029,123 bp) and *Lacticaseibacillus paracasei* ZFM54 shows 2,945 contigs (3,015,887 bp). In addition to the difference in genome length, the two isolates also showed variability in coding sequences on forward strand, genomic scaffolds, coding sequences on reverse strand, GC content, and GC skew (**Figure 2a-b**).

**Figure 2.**
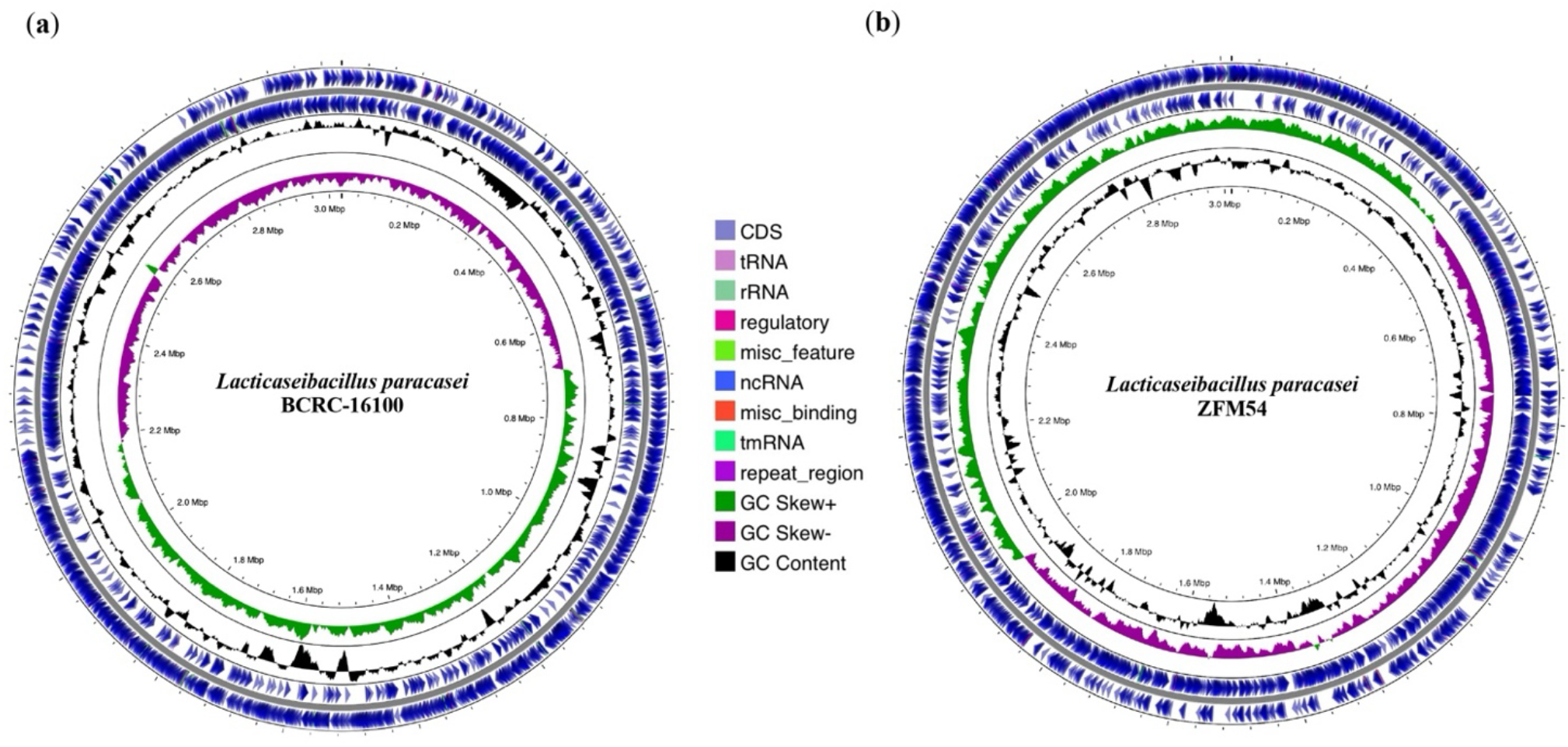
Whole genome sequence (WGS) assembly and analysis of *Lacticaseibacillus paracasei* BCRC-16100 and *Lacticaseibacillus paracasei* ZFM54. (**a**) Visualization of regions of interest in the whole genome of *L. paracasei* BCRC-16100 showing 2,885 contigs (3,029,123 bp) and (**b**) Visualization of regions of interest in the whole genome of *L. paracasei* ZFM54 showing 2,945 contigs (3,015,887 bp). Circles from the outside to the center illustrate the following characteristics: (1) coding sequences on forward strand, (5) genomic scaffolds, (2) coding sequences on reverse strand, (3) GC content, and (4) GC skew. Genetic maps were generated using Proksee v1.0.0a6.

The tolerance of *Lacticaseibacillus paracasei* BCRC-16100 and *Lacticaseibacillus paracasei* ZFM54 were tested in ranges of pH (3-7), alcohol concentration (5%-10%), and temperature (25°C-50°C) in 24h incubation (**Figure 3a-c**). **Figure 3a** shows that *L. paracasei* ZFM54 grows in a wider range from 35°C-45°C with 45°C as the optimal temperature. The optimal temperature for growth *L. paracasei* BCRC-16100 is the same but there is a significant decline in growth for any deviation in the temperature (±1°C). However, it has been previously reported that the growth for different *Lactobacillus* spp. (*rhamnosus, paracasei, reuteri, plantarum* and *pentosus*) is optimal at 37°C (Śliżewska and Chlebicz-Wójcik, 2020). This can be explained in the differences in the temperature points observed where the previous study observed growth in 4°C, 20°C, 30°C, 37°C, 44, and 55°C. In the different concentrations of alcohol tested (5%, 10%, 15%, and 20%), growth for the two isolates were the same suggesting the ability of the isolates to survive when incorporated in functional food products or drugs with alcohol content ranging from 5°C - 20% (**Figure 3b**). For both probiotic isolates, the growth at pH 5 and pH 7 were significantly higher than pH 3 and pH 4 (**Figure 3c**). This is consistent with the result of a previous study indicating that optimal pH for the growth of *Lactobacillus* spp. (*rhamnosus, paracasei, reuteri, plantarum* and *pentosus*) is 5.5 to 6.5 which may vary depending on the strain [43].

**Figure 3.**
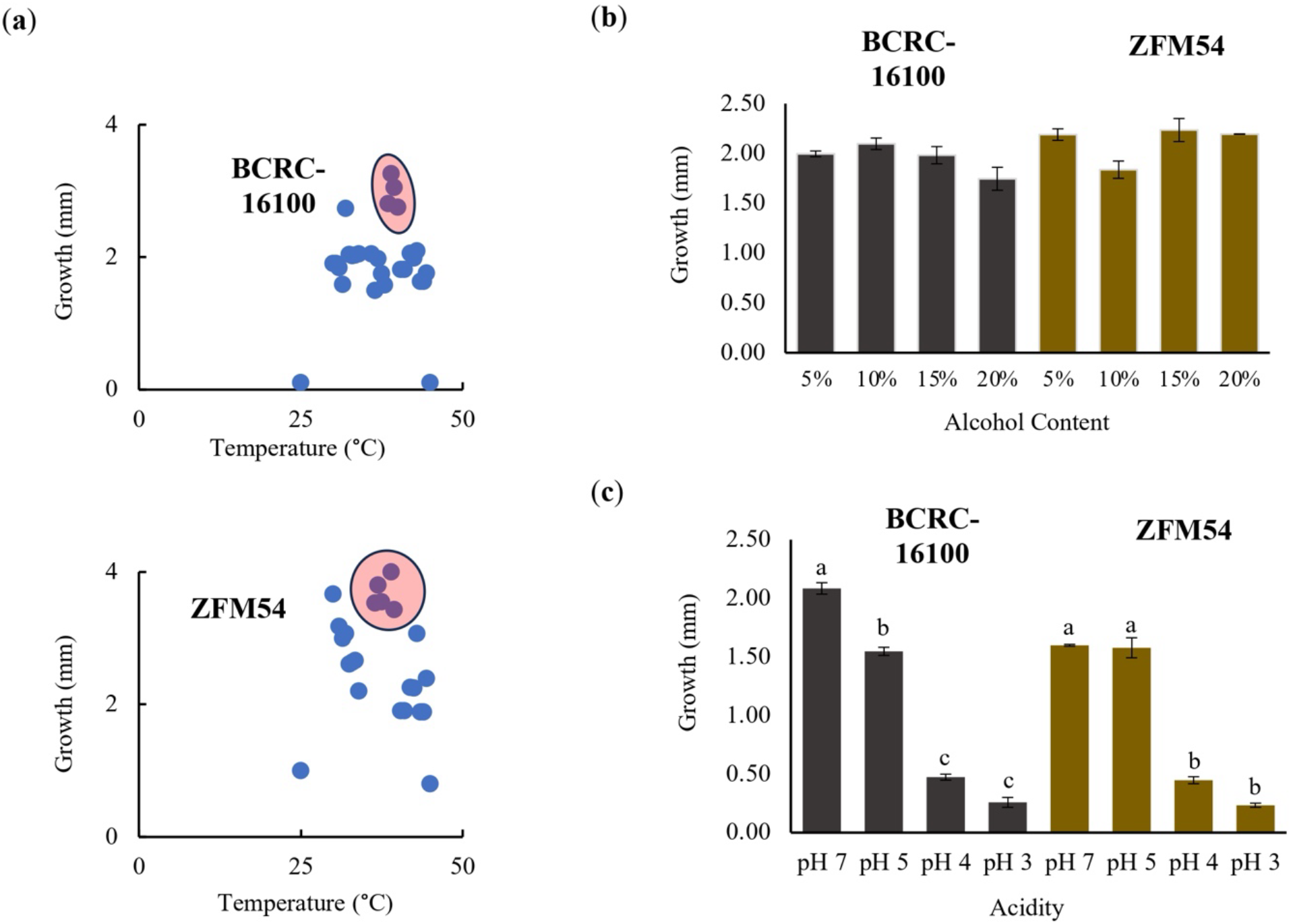
Tolerance of lactic acid bacterial isolates from nipa sap, *Lacticaseibacillus paracasei* BCRC-16100 and *Lacticaseibacillus paracasei* ZFM54. *L. paracasei* ZFM54 has wider range of tolerance across (**a**) temperature and (**b**) presence of alcohol while (**c**) *L. paracasei* BCRC-16100 performs better in a range of pH. Means within each bar (comparison within the same strain) having the same letter are not significantly different in Tukey’s multiple comparison at α = 0.05. and significant for condition as source of variation at *p* ≤ 0.01. Error bars represent standard error within 3 biological replicates of each type of condition.

The differences in the frequency of the stress tolerance genes and variability in the type and number of transcription factors (TFs) binding to cis-regulatory elements offers an explanation in differential or similar growth response of the two isolates in a particular stressful condition (**Figure 4a-c**). The growth of *L. paracasei* ZFM54 is less influenced by temperature and higher frequency of TFs that may potentially bind to its promoter region to a thermoregulatory gene *dnaK* can play a significant role in this tolerance. A previous study has shown that in known thermotolerant *Bacillus pumilus, dnaK* is highly expressed [44]. Although it cannot be ruled out that there might be differences in *dnaK* of different species and strains and other genes present in the genome of the two isolates play synergistic or antagonistic roles in tolerance to certain temperatures.

**Figure 4.**
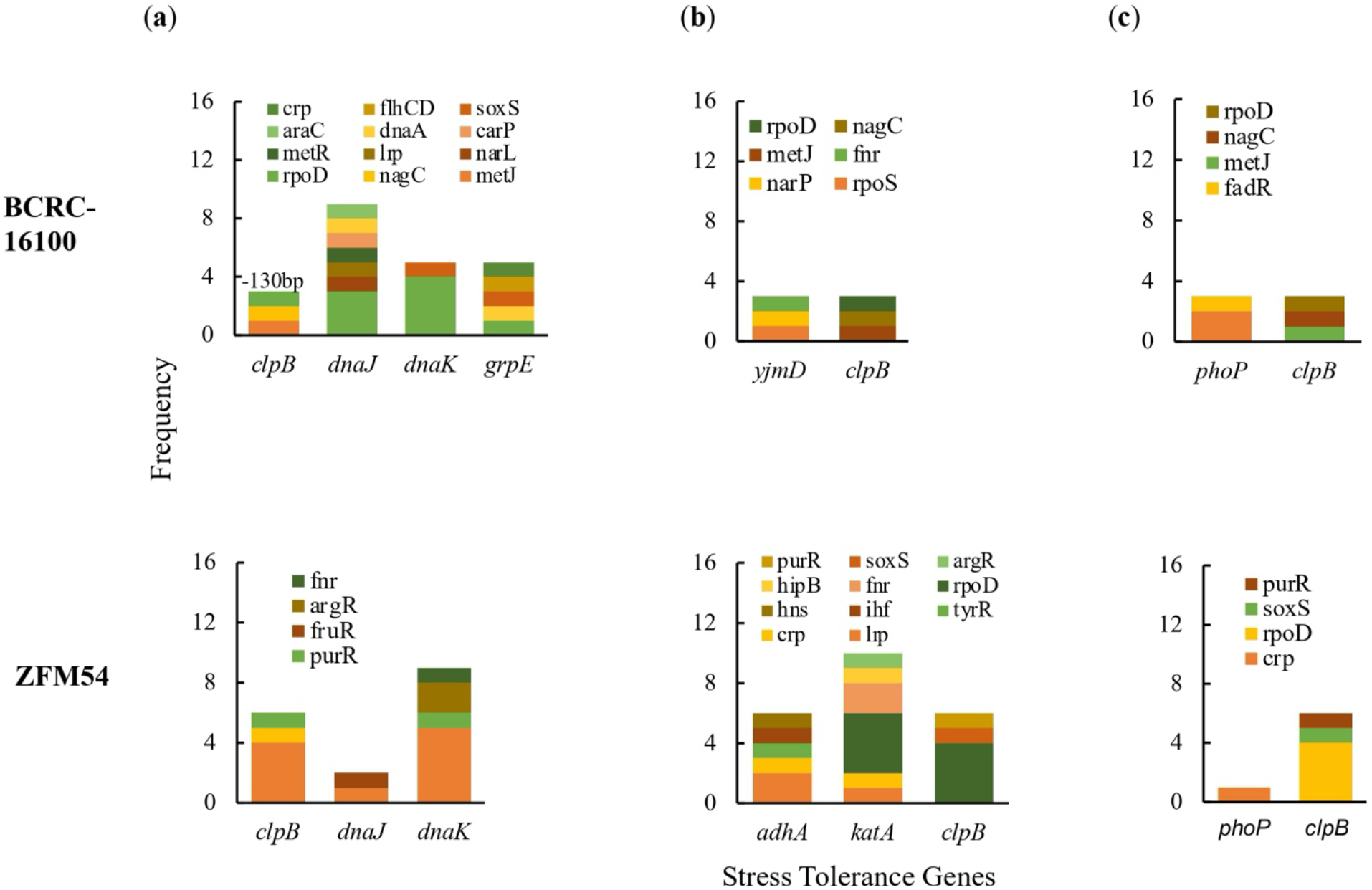
Promoter analysis of genes associated to (**a**) thermoregulation, (**b**) alcohol tolerance, and (**c**) pH response in *Lacticaseibacillus paracasei* BCRC-16100 and *Lacticaseibacillus paracasei* ZFM54. There is differential frequency in the number of tolerance genes and transcription factors binding to each gene responsible for stress response. Colors in the bars indicate the number of each type of transcription factors (TF) binding to a stress tolerance gene. Complete list of stress tolerance genes with the cis-regulatory elements where TFs bind is shown in Supporting Figure 11.

### 3.3. Antibiotic Resistance

The antibiotic resistance of the two probiotic isolates showed similar pattern in six (6) types of antibiotics both at 24h and 48h post-treatment (**Figure 5a-b**). In 24h, *Lacticaseibacillus paracasei* BCRC-16100 and *Lacticaseibacillus paracasei* ZFM54, showed complete resistance to clindamycin, vancomycin, gentamicin, ofloxacin, erythromycin, and streptomycin. However, at 48h post-treatment, five antibiotics (clindamycin, gentamycin, ofloxacin, erythromycin, and streptomycin) showed inhibition in the growth of two isolates except for vancomycin (**Figure 5a**). Differences in the zone of inhibition (mm) were significantly different (p=0.0005, α=0.05) in various antibiotics for each of the two isolates (**Figure 5b**). Higher zone of inhibition was observed in erythromycin while no inhibition was observed in vancomycin for the two isolates. The result on disk diffusion is corroborated by the colony forming unit quantity (CFU/ml) when the two isolates were independently grown in liquid broth infused with antibiotics for 48h. The CFU for both isolates were significantly higher in vancomycin (BCRC-16100=6.330413773, and ZFM54=5.252853031) suggesting resistance in this antibiotic that also support results of the disk diffusion analysis. These analyses indicate differential resistance of *L. paracasei* BCRC-16100 and *L. paracasei* ZFM54 depending on the type of antibiotic and time of exposure. Moreover, it also suggests the preponderance of antibiotic resistance genes containing active cis-regulatory elements in these isolates promoting survivability in antibacterial agents.

**Figure 5.**
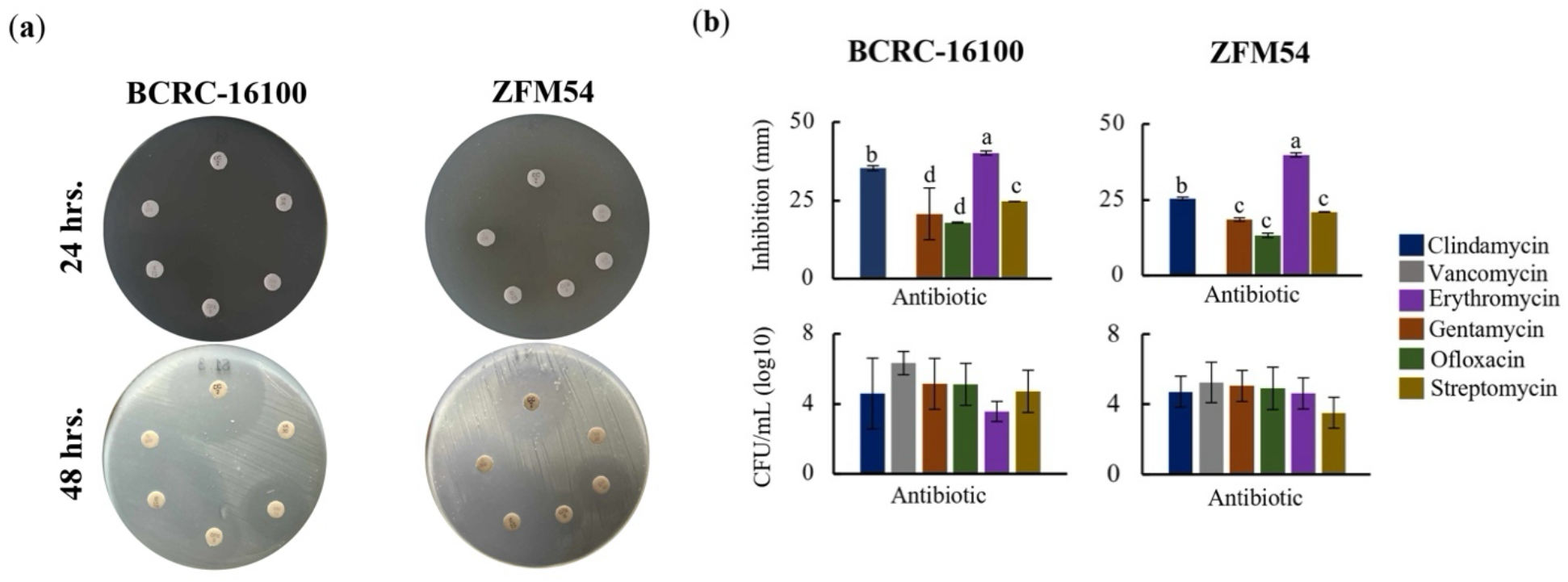
Higher antibiotic resistance in *Lacticaseibacillus paracasei* ZFM54 compared to *Lacticaseibacillus paracasei* BCRC-16100 suggested by higher growth in six (6) types of antibiotics and expression of antibiotic resistant (AbR) genes. (**a**) disk diffusion assay, zone of inhibition in antibiotic disks and (**b**) and colony forming units (CFU) of the two isolates in different antibiotics at 12h. From top clockwise in (a) the antibiotics are clindamycin, vancomycin, gentamycin, ofloxacin, erythromycin, and streptomycin. Means within each bar (comparison within the same strain) having the same letter are not significantly different in Tukey’s multiple comparison at α = 0.05. and significant for type of antibiotic as source of variation at *p* ≤ 0.01. Error bars represent standard error within 3 biological replicates of each type of condition.

### 3.4. Genomic Analysis of Antibiotic Resistant (AbR) Genes

AbR genes are present in both probiotic isolates which differ in terms of frequency, activity, and genomic, and amino acid structure (**Figure 6a-d**). There are five (*bmr3, stp, lmrA, emrY, yheI*) and nine (*stp, yheI, yheH, bmr3, tetA_1, lmrA, tetA_2, tetO, marR*) AbR genes in *Lacticaseibacillus paracasei* BCRC-16100 and *Lacticaseibacillus paracasei* ZFM54, respectively (**Figure 6a**). Briefly, there are two antibiotic resistant mechanisms (transcriptional regulation and transporter activity) in *Lacticaseibacillus paracasei* BCRC-16100 while there are three in *Lacticaseibacillus paracasei* ZFM54 suggesting that the later can have a wider range of resistance to antibiotics (transcriptional regulation, transporter activity, and efflux system). In *Lacticaseibacillus paracasei* BCRC-16100, the antibiotic resistant genes are mostly responsible in encoding transporter protein subunits (*bmr3, stp, emrY, yheI*) and one is a transcriptional repressor (*lmrA*). The two isolates share four AbR genes namely *bmr3, stp, lmrA, yheI* and unique AbR genes in *Lacticaseibacillus paracasei* ZFM54 include *yheH, tetA_1, tetA_2, tetO*, and *marR* where three are efflux pumps (*tetA_1, tetA_2*, and *tetO*), one encodes a transcriptional regulator (*marR*) and another that encodes a transporter protein subunit (*yheH*).

**Figure 6.**
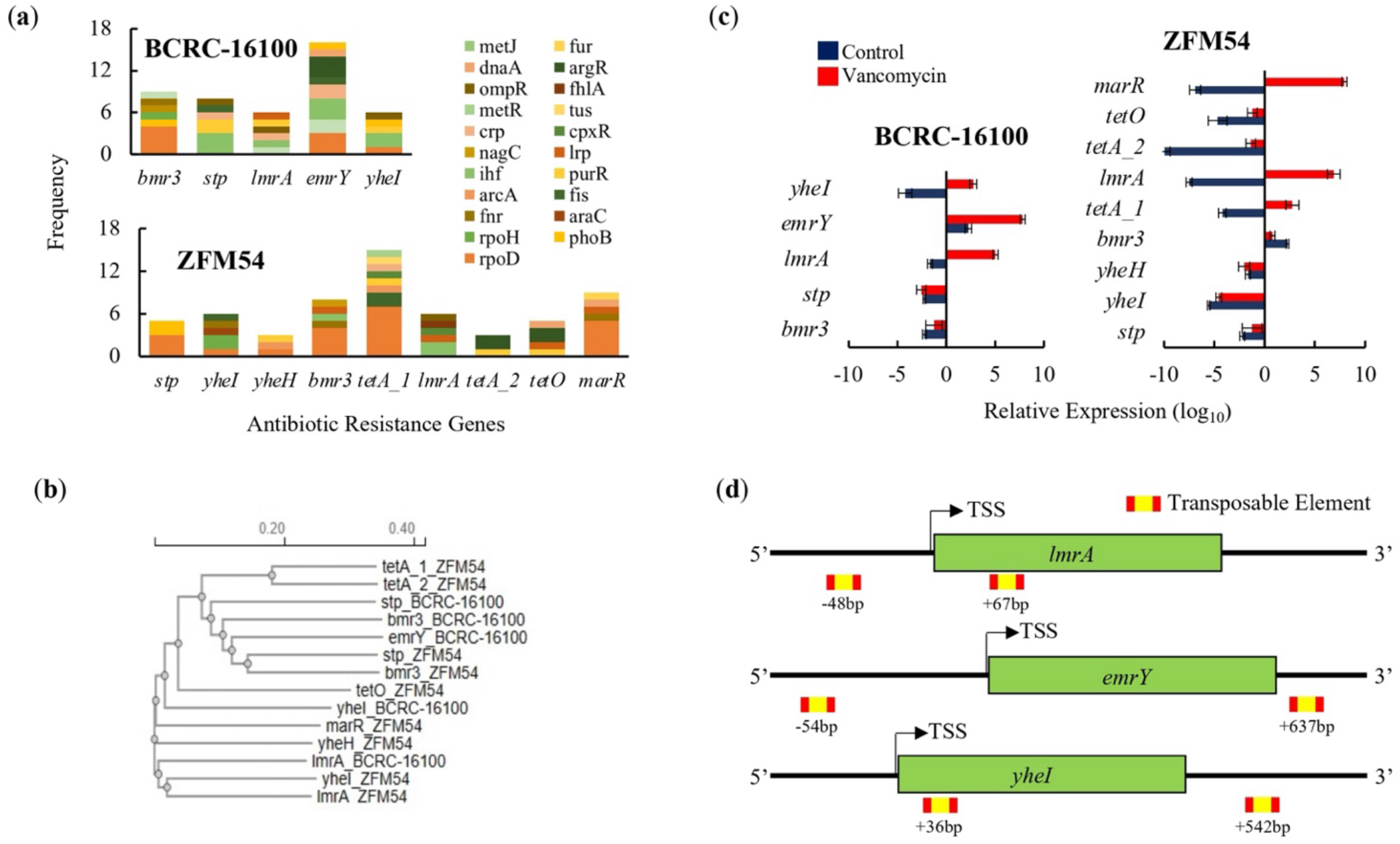
Promoter, phylogeny, gene expression, and transposable elements of antibiotic resistant (AbR) genes in *Lacticaseibacillus paracasei* BCRC-16100 and *Lacticaseibacillus paracasei* ZFM54. (**a**) AbR genes and frequency of each type of transcription factor (TF) binding in the promoter region (1000bp) in every gene. More resistant genes are found in the genome of *L. paracasei* ZFM54 and the frequency of TFs that can bind to each of the AbR genes promoter region vary. (**b**) Phylogeny of AbR genes showing differential clustering of some similar genes (*stp* and *bmr3*) shared by the two isolates. (**c**) Expression of AbR genes in the two isolates. (**d**) Transposable elements located in the upstream (5’) and downstream (3’) regions (500bp) of AbR genes. Colors in the bars indicate the number of each type of TF binding to a stress tolerance gene. Complete list of AbR genes with the cis-regulatory elements where TFs bind is shown in Supporting Figure 10. Relative expression is against the internal control (*16s rRNA*) is shown. Transcription Start Site (TSS). Error bars represent standard error within 3 biological replicates.

Although four AbR genes are shared between *Lacticaseibacillus paracasei* BCRC-16100 and *Lacticaseibacillus paracasei* ZFM54, analysis of the homology of these four genes relative to each other including the rest of AbR genes indicate that there are differences. Among the four shared AbR genes in the two isolates, *yheI* and *stp* are the most diverged as suggested by the distance in their cluster or clades in the phylogenetic analysis (**Figure 6b**). Given the overlapping and diversity of similar AbR genes in the two isolates, genes responsible for antibiotic resistance were tested for differential gene expression analysis in the presence in vancomycin (**Figure 6c**), an antibiotic where both isolates showed resistance (**Figure 5a**). For both isolates, AbR genes encoding transcriptional regulators (*lmrA and marR*) are upregulated in vancomycin treatment. This may suggest a similar mechanism for vancomycin resistance in *Enterococci* where vancomycin activates *VanR* which then cascades to the activation of other AbR genes [45]. Other AbR genes that are significantly upregulated in vancomycin are *emrY* and *yheI* which are encoding for transporters protein subunits and *tetA_1* encoding for an efflux pump. Among the AbR genes, only *emrY* has the highest probability to be transposed laterally due to the presence of transposable elements in the - 54bp (5’) and +637bp (3’) of the gene (**Figure 6d**).

### 3.5. Carbohydrate Hydrolytic Activity

The two isolates were analyzed for ability to hydrolyze carbohydrates and TF binding to the promoter and expression of genes responsible for hydrolysis (**Figure 7**). In a starch assay, both isolates showed hydrolytic activity qualitatively (**Figure 7a**) and quantitatively (**Figure 7b**). There were multiple genes found in the WGS which may be responsible for the carbohydrate hydrolytic activity having 8 and 11, in *Lacticaseibacillus paracasei* BCRC-16100 and *Lacticaseibacillus paracasei* ZFM54, respectively (**Figure 7c**). Notably, promoter analysis showed more TF binding to *bga, lacG* and *bglA* suggesting the importance of the three genes in carbohydrate analysis. However, gene expression analysis revealed that during carbohydrate hydrolysis, *rafA, malL*, and *bglC* are the genes that are upregulated. It can be argued that despite the presence of more cis regulatory elements in some carbohydrate hydrolysis genes, the observed lower expression can be explained as a result of positional effect of where the TF bind which was previously described to influence gene activity [46].

**Figure 7.**
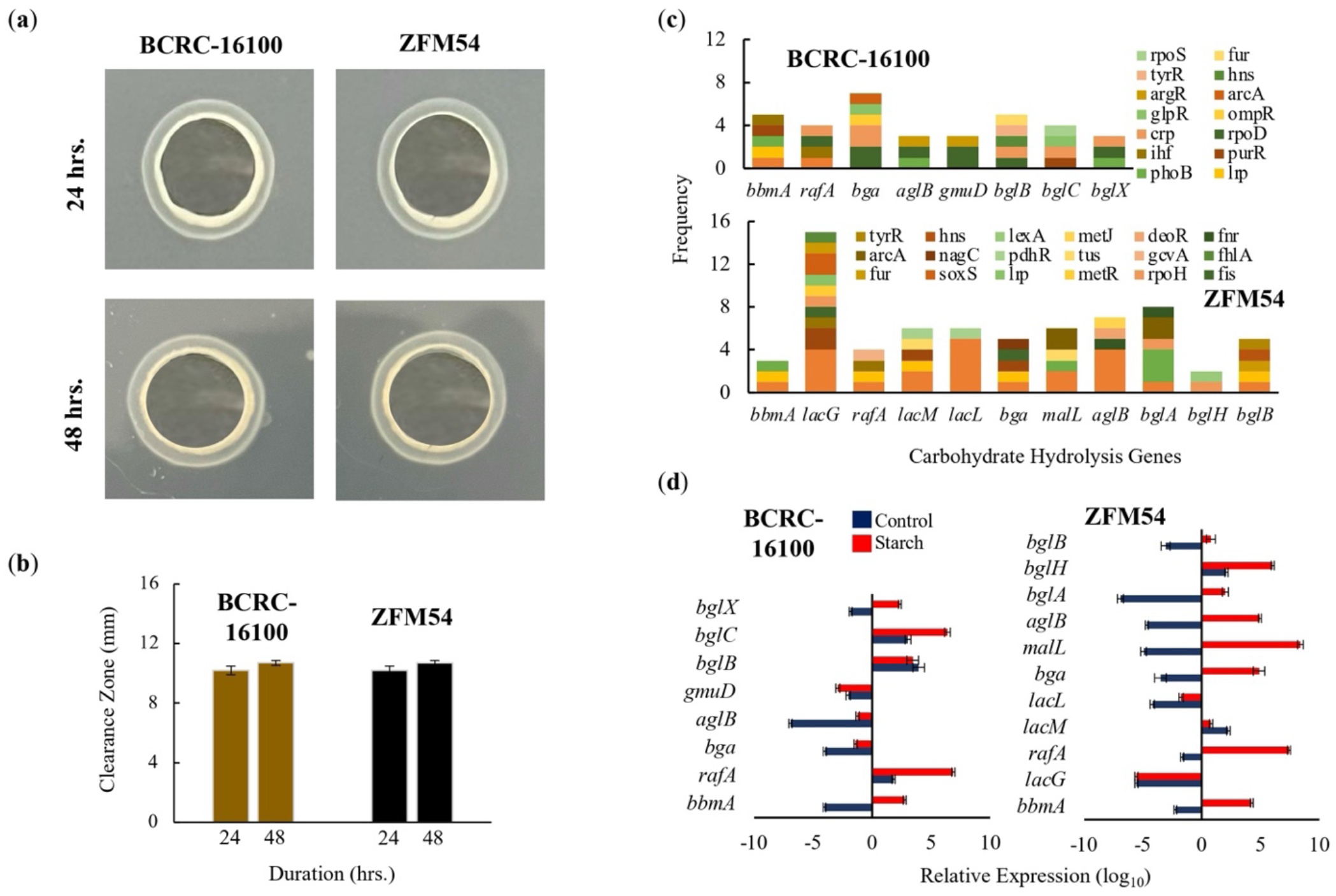
Carbohydrate hydrolytic activity of *Lacticaseibacillus paracasei* BCRC-16100 and *Lacticaseibacillus paracasei* ZFM54. (**a**) Amylase activities of BCRC-16100 and ZFM54 presented by zone of clearance, (**b**) the quantitative zone of clearance expressed in mm, (**c**) promoter analysis of carbohydrate hydrolysis genes, and (**d**) expression of amylases genes. Colors in the bars indicate the number of each type of transcription factors (TF) binding to a stress tolerance gene. Relative expression is against the internal control (*16s rRNA*) is shown. Error bars represent standard error within 3 biological replicates.

## 4. DISCUSSION

Probiotics has been widely used for numerous health benefits such as improved gut microflora, enhance digestion of nutrients and reduction in the risk of various diseases [47-48]. Despite the known positive health effects, it is also essential to ensure the (1) flexibility of probiotic isolates in different conditions to allow incorporation in drugs and functional food, (2) absence of transposable antibiotic resistance genes, (3) and bioactivity of any proposed benefit such as carbohydrate digestion. Probiotics can be isolated from animal and dairy sources where novel or similar genotypes obtained from a different source can have different characteristics [49-51]. However, efforts to isolate probiotics from plant sources are very limited [52].

There are multiple indigenous plants in northern Philippines that can be instrumental as sources of probiotics due to their abundance in the region such as nipa. Two strains of the same species of probiotics can be isolated from this source namely *Lacticaseibacillus paracasei* BCRC-16100 and *Lacticaseibacillus paracasei* ZFM54. The genus of the two strains is classified as probiotics but there is very little to know understanding on their health benefits, tolerance to stress and safety. Moreover, the species of the two strains are also recognized as probiotics only known for ameliorating allergic airway, antimicrobial activity, enhancement of intestinal microbiota and stress modulator, among others [53-55]. The intrinsic heterogeneity across strains of *L. paracasei* makes the species a strong group for the selection of probiotics isolates for health benefits and development of drugs or functional food for the abovementioned purpose [56]. Although in recent years much effort has been made to study this, the answer is not conclusive, and there remains much to be elucidated.

As described above, the health-promoting traits from the species *L. paracasei* as probiotics has not been extensively explored and the strains isolated from nipa sap, *L. paracasei* BCRC-16100 and *L. paracasei* ZFM54, has not been characterized for its contribution to overall health, compatibility for drug and functional food development and absence of transposable antibiotic resistance elements.

To assess the potential of *L. paracasei* BCRC-16100 and *L*.*s paracasei* ZFM54 as probiotic for commercial use, we carried out whole genome sequence, gene expression and biochemical analyses. There are several genes that confer thermotolerance in both isolates which allows growth in a wide range of temperature where *dnaK* and *dnaJ* are likely responsible for this trait these genes show the highest potential to be regulated due to the association of promoter regions to multiple transcription factors (TFs) (**Figure 4a**). Both *dnaK* and *dnaJ* belong to a chaperone system in microbes known to confer wider tolerance to temperatures [44,57]. The two isolates grow optimally at pH 5-7, however, promoter analysis of genes responsible for this trait only suggested *clpB* as may be responsible in differential growth in pH. The gene *clpB* is highly conserved in bacteria to provide tolerance to oxidative stress, nutrient starvation, and low pH [58-60]. In addition to flexibility to stressful conditions, we also explored AbR genes in each strain with the corresponding molecular mechanisms and transposability. The two strains were highly resistant to vancomycin and gene expression analysis suggests four genes conferring this resistance namely *lmrA, marR, emrY* and *yheI* and mechanisms of resistance include efflux, transcriptional control, and regulation of transport (**Figure 6**). Moreover, these AbR genes possess transposable elements in the upstream (5’) and downstream (3’) region. This is a consideration in using these strains as probiotics to avoid potential lateral transfer of AbR genes to pathogenic microbes [61]. Lastly, high rates of available genes that encode carbohydrate active enzymes are detected in the genome data, which correlated the enzymatic hydrolytic activities evident *in-vitro*. Among the identified genes, *rafA, malL*, and *bglC*, are most likely to be responsible to the carbohydrate hydrolytic activity of *L. paracasei* BCRC-16100 and *L. paracasei* ZFM54 based on the promoter analysis of genes (**Figure 7**). Bacteria that colonize the intestine collectively possesses a large repertoire of degradative enzymes and metabolic capabilities. To promote the development of human gastrointestinal nutrition and health is to regulate the host mucosal and balance intestinal microflora.

In conclusion, *L. paracasei* BCRC-16100 and *L. paracasei* ZFM54 can arguably be utilized as probiotics that can be incorporated in drugs and functional food to promote digestion of carbohydrates in food products with low digestibility with careful consideration of the antibiotic resistance in these strains.

## Supplementary Materials

Table S1: Primers for PCR and qPCR and sequencing *Lacticaseibacillus paracasei* BCRC-16100 and *Lacticaseibacillus paracasei* ZFM54; Table S2: Analysis of Variance for alcohol tolerance of *Lacticaseibacillus paracasei* BCRC-16100; Table S3: Analysis of Variance for alcohol tolerance of *Lacticaseibacillus paracasei* ZFM54; Table S4: Analysis of Variance for pH tolerance of *Lacticaseibacillus paracasei* BCRC-16100; Table S5: Analysis of Variance for pH tolerance of *Lacticaseibacillus paracasei* ZFM54; Table S6: Analysis of Variance for inhibition zone of *Lacticaseibacillus paracasei* BCRC-16100 in five antibiotics (clindamycin, vancomycin, gentamycin, ofloxacin, erythromycin, and streptomycin); Table S7: Analysis of Variance for inhibition zone of *Lacticaseibacillus paracasei* ZFM54 in five antibiotics (clindamycin, vancomycin, gentamycin, ofloxacin, erythromycin, and streptomycin); Table S8: Analysis of Variance for CFU/mL of *Lacticaseibacillus paracasei* BCRC-16100 in five antibiotics (clindamycin, vancomycin, gentamycin, ofloxacin, erythromycin, and streptomycin); Table S9: Analysis of Variance for CFU/mL of *Lacticaseibacillus paracasei* ZFM54 in five antibiotics (clindamycin, vancomycin, gentamycin, ofloxacin, erythromycin, and streptomycin); Table S10: Antibiotic Resistant (AbR) Genes identified in the genome sequences of *Lacticaseibacillus paracasei* BCRC-16100 and *Lacticaseibacillus paracasei* ZFM54, associated with their corresponding transcription factors (TF) and respective binding site scores; and Table S11: Name of the isolates and results of the identification.

## Author Contributions

Conceptualization, P.J.I.G.; data curation and formal analysis, P.J.I.G..; funding acquisition, S.C.A. and D.S.B.; investigation, M.T.A., J.Z.P.F., S.M.B., A.J.R., M.P.O., J.F.C., and A.D.; project administration, S.C.A., D.S.B., and P.J.I.G.; software, R.J.J.P. and A.G.B.P.; supervision, P.J.I.G.; validation, P.J.I.G.; writing – original draft, P.J.I.G., J.Z.P.F, and A.G.B.P.; writing – review and editing, P.J.I.G., J.Z.P.F., and M.T.A. All authors have read and agreed to the published version of the manuscript.

## Data Availability Statement

The complete genome sequence of the isolates investigated in this study can be accessed in the NCBI GenBank Database, with accession numbers, NZ_CP086132 and NZ_CP032637.

## Acknowledgments

This project was supported by the Commission on Higher Education (CHED) in the Philippines with grant number LAKAS 2022-005. We thank Policarpio B. Joves for providing a microbiology laboratory.

## Conflicts of Interest/Disclaimer

The authors declare no conflicts of interest.

